# Analysis of Circuits for Dosage Control in Microbial Populations

**DOI:** 10.1101/2020.12.18.423556

**Authors:** Sophie J. Walton, Samuel E. Clamons, Richard M. Murray

## Abstract

Designing genetic circuits to control the behaviors of microbial populations is an ongoing challenge in synthetic biology. Here we analyze circuits which implement dosage control by controlling levels of a global signal in a microbial population in face of varying cell density, growth rate, and environmental dilution. We utilize the Lux quorum sensing system to implement dosage control circuits, and we analyze the dynamics of circuits using both simplified analytical analysis and *in silico* simulations. We demonstrate that strong negative feedback through inhibiting LuxI synthase expression along with AiiA degradase activity results in circuits with fast response times and robustness to cell density and dilution rate. We find that degradase activity yields robustness to variations in population density for large population sizes, while negative feedback to synthase production decreases sensitivity to dilution rates.

## Introduction

A challenge in synthetic biology is to engineer microbial consortia to perform coordinated tasks as a population with behaviors and functions that are not present or possible in single cells [1, 2]. In order to engineer such microbial consortium, we need to develop population level controllers that exhibit robustness to environmental disturbances [1]. Despite examples of circuits that yield robustness on a single cell level [3–5], there has been less focus on creating circuits that are able to control levels of a global species that is produced by a collection of cells. This is difficult, for it requires cells to be able to communicate with one another in order to coordinate their behavior. Quorum sensing systems in bacteria can be used to communicate between cells and adjust cell responses [6]. Since quorum sensing molecules can convey information about population density, these molecules can be utilized for designing synthetic circuits for population level control [7]. For example, N-Acyl homoserine lactones (AHLs) have been used to implement synchronized genetic oscillators [8, 9], control cell density [10], and maintain robustness across heterogeneous populations [11–14]. These efforts demonstrate the power of AHLs to be used as sensors and controllers in order to coordinate population level behaviors in synthetic circuits.

One potential application of engineered microbial populations would be to release constant doses of a drug into the gut of a patient. The microbial population could sense the conditions of their external environment and together adjust production of the drug such that a patient consistently receives a constant dose. In order for a patient to receive an appropriate dose, the levels of drug in the patient’s gut need to be robust to environmental factors. For example, changing nutrient concentrations could alter the growth rate of the population and cause the population size to fluctuate. We thus need a population level controller that is robust to such disturbances, and such a circuit would implement dosage control.

We explore simple circuits to implement dosage control of AHLs using the Lux quorum sensing system. Previous work demonstrated that a closed loop circuit in which AHLs activated expression of AiiA degradases, molecules that degrade AHLs, the global concentration of AHLs was more robust to changing population density [15]. However, this circuit was not robust in low population densities, and did not explore whether or not this circuit was robust to varying growth rates and reactor dilution rates. Additionally, there have been studies on the utilities of LuxR repressive quorum sensing systems for reducing variability of protein expression in cellular populations [13, 14], and these circuits can also be used to reduce variability of AHL expression due to varied population size. We expand on this work and evaluate various dosage control architectures that are implemented using species from the Lux quorum sensing system. We used simplified analytical analysis to gain intuition into the roles of AHL degradases and Lux repressive quorum sensing systems. Through simulations we demonstrated that AHL degradation leads to robustness to population density in dense populations, and repression of synthase molecules can reduce sensitivity to dilution rates. We found that layered feedback through a repressive quorum sensing system and a secondary repressor can be utilized to increase robustness to population density. Additionally, we evaluate how tunable each circuit is, and we examine the trajectories of each circuit through evaluating properties such as overshoot and settling times.

## Results

### Analytical Analysis of Simplified Dosage Control Circuits

In the following analysis we model the dynamics of AHLs in a bioreactor with a homogeneous population of cells at optical density (OD). OD is unitless, and here we use it as a proxy for cell density to be consistent with [15]. For *Escherichia coli* 1 OD ≈ 10^8^ cells/mL. The cells are all growing at rate *γ*_*g*_ and media containing cells, waste, and global species flows out of the reactor at rate *γ*_*r*_. We assume that AHL diffusion in and out of cells is fast, and thus the external and internal concentration of the shared signal is equal [10, 15, 16]. Additionally, AHLs degrade in the reactor medium at a rate *γ*_*H*_ [17]. Our goal is to find a circuit such that the concentration of AHLs in the reactor is robust to changes in population density and dilution due to growth (*γ*_*g*_) and reactor flow (*γ*_*r*_). We use the Lux quorum sensing system as our model for dosage control to be consistent with previous work on dosage control [15].

We explore the dynamics of six circuits under simplifying assumptions in order to gain intuition in the roles of AHL degradases and AHL genetic feedback. Even though some of these assumptions are known to be inconsistent with the system we are studying, these assumptions make analysis more amenable in order to gain intuition for the roles of different parameters and species in each circuit before conducting full modeling. We assume that *N*_*v*_ (OD) is at steady state, and thus we trear *N*_*v*_ as a parameter rather than a model variable. In our simplified analysis we solve for steady states of certain systems with the following assumptions about the dynamics of certain model species:

1. unless indicated, all Hill coefficients are 1;
2. for activating Hill functions of the form 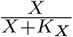 we assume that *K*_*X*_ ≫ *X* such that 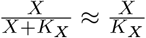 (which is equivalent to assuminng that enzymes are not saturated) [15];
3. we assume that leak parameters are negligible.

We first analyzed an open loop controller and a closed loop controller that consist of only AHLs (*H*) and LuxI synthase molecules (*I*) (Fig 1A). LuxI synthesizes AHLs from S-adenosylmethionine, which is has a high intracellular concentration [6, 18, 19]. The open loop controller is described in [15], and it consists of LuxI synthase (*I*) being produced at rate *α*_*I*_ and diluted due to growth at rate *γ*_*g*_. We assume that *I* is tagged with an LVA degradation tag such that they are degraded at a rate *γ*_*d*_. LuxI produces AHLs at rate proportional to an activating hill function of *I*. The dynamics of this open loop circuit can be modeled with the following ODEs:

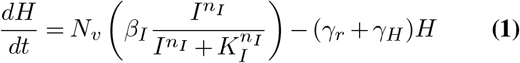

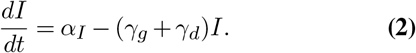

With the previously described simplifying assumptions we can solve for the steady state of this system:

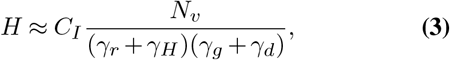

where 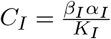. As noted in [15], this circuit is not robust against changes in *N*_*v*_ or dilution rate in any regime of *N*_*v*_.

**Fig. 1.**
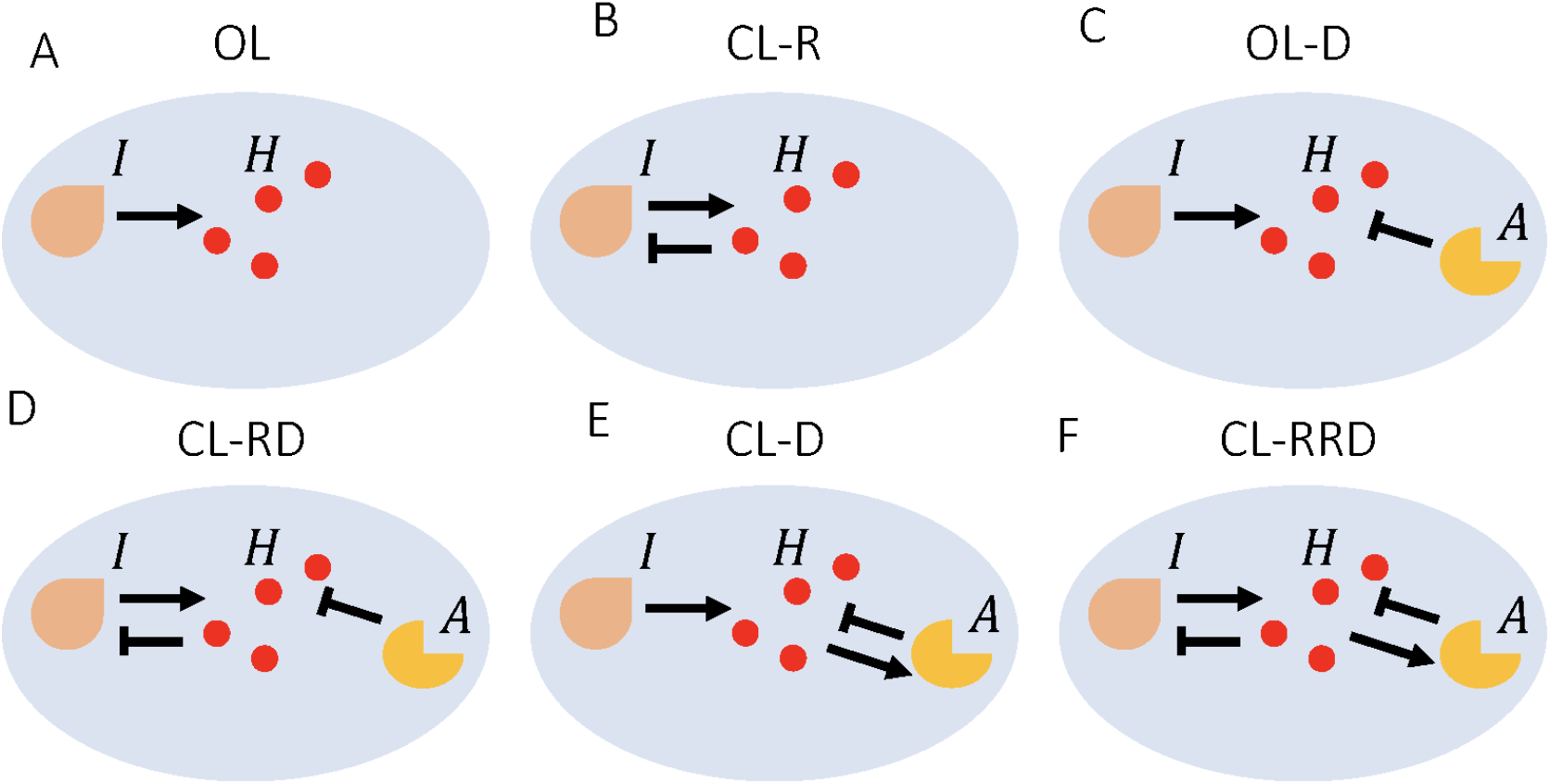
Diagrams of open loop (A, C) and closed loop (B,D,E,F) circuit architectures described. *H* represents AHLs, *I* respresents LuxI synthase, *A* represents AiiA degradase.

In the closed loop circuit we have a repressive quorum sensing system controlling the dynamics of *I*, which has been described and analyzed previously in [13, 14]. This circuit is thus called Closed Loop Repression (CL-R) (Fig. 1B). AHLs, *H*, control transcription of proteins by binding to a transcription factor, LuxR, which regulates expression of genes under a pLux promoter. We do not model the dynamcis of the LuxR protein, for we assume that it is constantly expressed and abundant [8, 9, 15]. In the CL-R circuit we assume that *I* under the control of a promoter that is repressed by LuxR [20].

This can be modeled with equation (1) for the dynamics of *H* and the following equation for the dynamics of *I*:

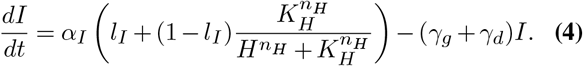

First we solve for the steady state through assuming that 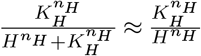 where *n*_*H*_ ∈ ℝ^+^, which models the regime where we have excess of *H*. Under this assumption the steady state level of *H* is approximately:

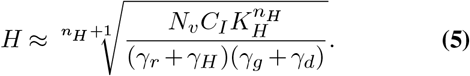

We see that this is sensitive to dilution rate and population density. In the above analysis we assumed that *H* is in abundance with respect to *K*_*H*_, but the steady state level of *H* changes if we instead assume that *H* ≈ *K*_*H*_. We care about this regime because this is where we enter the effective dynamic range of the repressor. For a general repressor *X*, whose dynamics are represented by 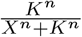 when *X* ≈ *K* the Hill repressing function can be approximated as

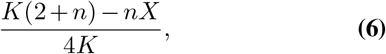

which was demonstrated in [13]. Thus if we assume that *H* ≈ *K*_*H*_ instead of *H* ≫ *K*_*H*_ we have that

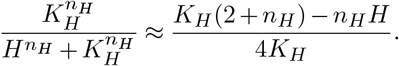

Under this assumption (along with the other previous ones) the steady state level of *H* is approximately

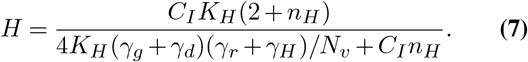

Here we see that as *N*_*v*_ → ∞ we have that 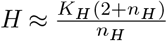 which is independent from both dilution rates and *N*_*v*_. Note that this only holds true in the regime where AHL (*H*) does not saturate the repressor controlling synthase production. This means that in order for *H* ≈ *K*_*H*_, we must have *n*_*H*_ ≫ 1.

From our above analysis we see that if we can tune the circuit such that *H* ≈ *K*_*H*_, then we can achieve robustness to dilution rate and population size with a simple repressive quorum sensing system. Previous dosage control analysis utilized AiiA degradases to achieve this robustness [15]. The dynamics of *H* for circuits utilizing degradases *A* are

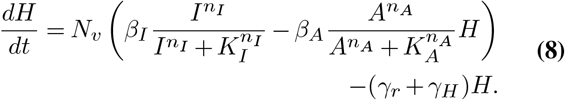

Consider an open loop circuit that contains AiiA degradases, which we call OL-D (Open Loop Degradation) (Fig. 1C). We assume that the degradase species (*A*) is tagged with an LVA degradation tag such that they are degraded at a rate *γ*_*d*_ [15], for this will increase the rate of *A* elimination to improve the speed of the circuit. Thus the dynamics of *A* are as follows:

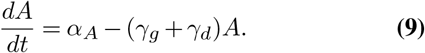

We use equation (2) for the synthase dynamics, and we solve for the steady state of *H* under our simplifying assumptions:

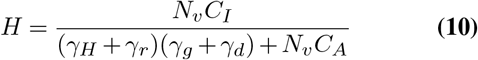

where 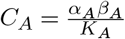. Note that *H* approaches 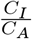 in the limit of large *N*_*v*_ relative to *γ*_*g*_, *γ*_*r*_. We thus see that this simple circuit can have desirable dosage control properties despite being an open loop controller. Although there is negative feedback from AHLs binding to degradases to trigger their own destruction, we call this an open loop circuit because the components of the circuit (*A, I*) are not subject to active regulation by AHLs (Fig. 1C). We can make this circuit closed-loop by regulating the expression of *I* as done in CLR (equation (4)). We call this circuit CL-RD (Closed Loop Repression Degradation) (Fig. 1D). We now use equation (4) to model the synthase dynamics. We first examine the case where

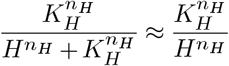

as done above. Under this assumption as well as the other previous ones the steady state level of *H* is approximately

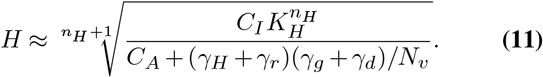

As *N*_*v*_ → ∞ we see that 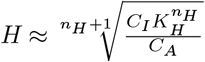, which is robust to both dilution rates and population size.

We next analyze the circuit in the case where *H* ≈ *K*_*H*_. Under this assumption (along with the other previous ones) the steady state level of *H* is approximately

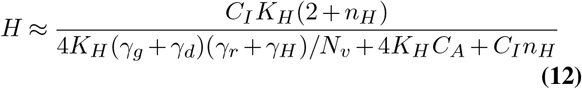

Here we see that as *N*_*v*_ → ∞ we have that 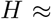 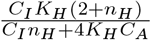 which is independent from both dilution rates and *N*_*v*_. Note that this steady state differs from the one obtained by assuming that *H* ≫ *K*_*H*_. Regardless of the working regime of the repressive system the steady state of *H* is independent of *γ*_*r*_, *γ*_*g*_, and *N*_*v*_ in the limit of large *N*_*v*_ and small *γ*_*r*_, *γ*_*g*_.

Next we examine circuits in which degradase (*A*) expression is regulated by *H*. We first outline the dynamics of a circuit described in [15] where synthase expression is unregulated but AHLs activate expression of the degradases (Fig. 1E). We call this circuit CL-D (Closed Loop Degradation). For simplicity we assume that LuxI is constituitively expressed at rate *α*_*I*_. We use equation (2) to model the dynamics of the synthase. We assume that *A* is under the control of a promoter that is activated by LuxR, and as done in [15] we do not model the dynamics of LuxR. Thus the following can be used to model the dynamics of *A*:

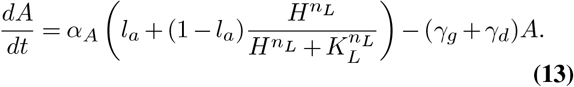

The authors of [15] showed that under the same simplifying assumptions made previously, the steady state of AHLs is

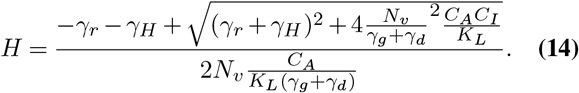

The above circuit is robust to dilution rates *γ*_*r*_, *γ*_*g*_ and OD (*N*_*v*_) in the limit of *N*_*v*_ → ∞. In this limit we have that 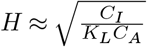 [15].

We analyze a circuit in which *I* is regulated as in equation (4) and *A* is regulated as in equation (13). We call this circuit CL-RRD (Closed Loop Repression Regulated Degradation) (Fig. 1F). We noted that in our other circuits in the limit that *N*_*V*_ → ∞ the steady state is independent of *N*_*v*_ and *γ*_*r*_. We assume that *N*_*v*_ ≫ 1 and *γ*_*r*_ ≪ 1 and attempt to solve for the steady state of the system. Since we assume that *N*_*v*_ ≫ 1 and *γ*_*r*_ ≪ 1 we have that

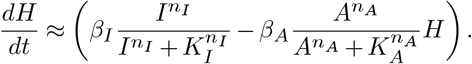

First, we solve for *H* when *H* ≫ *K*_*H*_ and obtain

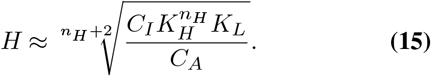

When we assume that *H* ≈ *K*_*H*_ instead, in the limit of *N*_*v*_ ≫ 1 and *γ*_*r*_ ≪ 1 we have that

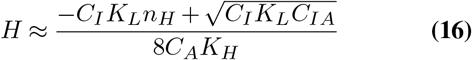

where 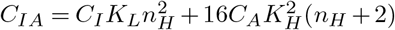.

From our simplified analysis we see that all closed loop circuits seem to have regimes where *H* will be robust to population density and dilution rate *γ*_*r*_ in the limit of large OD. However, we still do not know if this intuition will hold using parameters that are more representative of the components of the Lux quorum sensing system, and we still do not have an understanding of trajectories of these systems. In our next section we explore both of these issues through *in silico* simulations.

### Evaluation of Robustness to Population Density and Dilution

In the following sections we simulate the equations we derived in the previous section, but we use parameters derived from experiments characterizing the Lux quorum sensing system [15, 20]. Note that some of these parameters are not consistent with simplifying assumptions. As done in our previous analysis, we assume that *N*_*v*_ does not vary with time. We thus hold *γ*_*r*_ = *γ*_*g*_, which is a necessary condition for *N*_*v*_ to be constant and mimics conditions of a chemostat. We set the parameters governing enzyme dynamics (such as *K*_*d*_s for AiiA and LuxI) to be the same for all circuits, and we varied promoter strengths governing LuxI and AiiA expression such that most circuits were tuned to have a steady state of approximately 3 – 4 nM when *γ*_*g*_ = *γ*_*r*_ = 0.01 min^−1^ and *N*_*v*_ = 10.

We first characterized the dependence of the steady state of each circuit on population density and dilution rates (Fig. 2). Here we confirmed our previous prediction that the concentration of AHLs in circuits with degradases is robust to *N*_*v*_ as *N*_*v*_ becomes large (Fig. 2). Although CL-R performs better than OL, it does not display asymptotic robustness to *N*_*v*_ (Fig. 2A, B). This is because the repressor becomes saturated and the repressive quorum sensing system does not have high enough cooperativity (*n*_*H*_ ≈ 2). If *n*_*H*_ >> 1 and *α*_*I*_ ≈ *K*_*H*_, then the steady of *H* becomes more robust to population size and dilution rate (Fig. S1), but this assumption does not match experimental data [20]. On the other hand, every circuit that incorporates degradases displays robustness to *N*_*v*_ for high *N*_*v*_ as predicted by our simplified analysis (Fig. 2C – F). We also found the sensitivity of *H* to 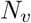 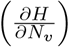 of each circuit (Fig. S2). All closed loop circuits and OL-D (but not OL) displayed decreasing sensitivity to *N*_*v*_ with increasing *N*_*v*_ (Fig. S2).

**Fig. 2.**
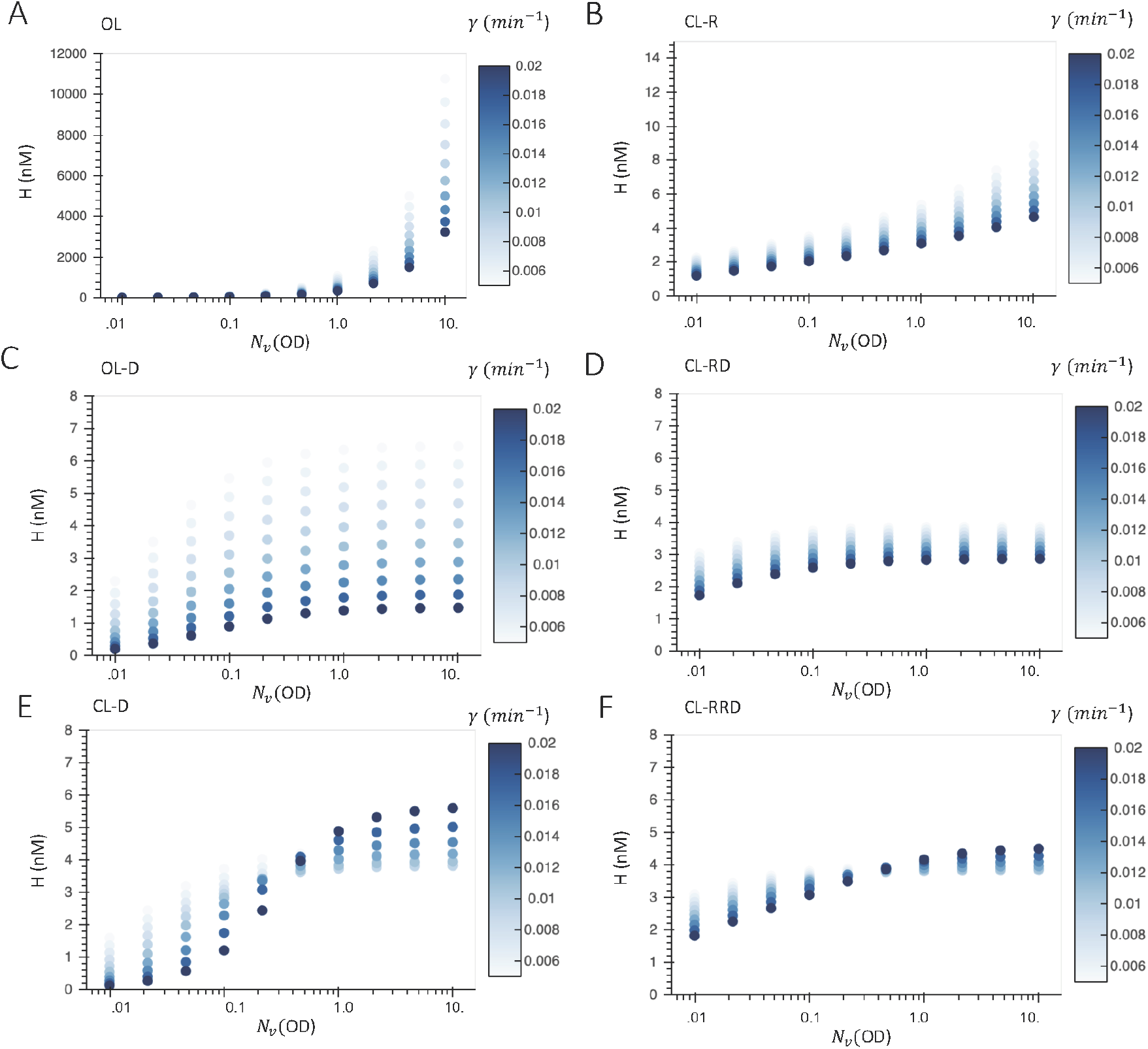
Steady state AHL concentrations for varying dilution rate(*γ* = *γ*_*r*_ = *γ*_*g*_) and OD (*N*_*v*_) for various dosage control architectures. Circuits were simulated for 1000 minutes, and the endpoint of the circuit was taken as the steady state of the circuit for a given set of parameters.

From our simplified analysis we saw that some of our circuits were predicted to be robust to dilution due to cell growth and reactor dilution in the limit of large *N*_*v*_, but this does not hold in our simulations. Circuits with regulated degradase expression (CL- RRD, CL-D) display minimum variation of *H* with dilution rates *γ* = *γ*_*r*_ = *γ*_*g*_ with intermediate *N*_*v*_ (Fig. 2E, F). Additionally, the magnitude of sensitivity to 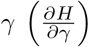 is minimized with intermediate *N*_*v*_ for intermediate *N*_*v*_ (Fig. S3E, F). Although sensitivity to *γ* plateaus for high *N*_*v*_ in circuits with unregulated degradase expression (OL-D, CL-RD), the sensitivity does not approach 0 (Fig. 2C, D; Fig. S3C, D). These results demonstrated that the dynamics of sensitivity to *γ* are mainly determined by the mode of degradase regulation, but we noted that negative feedback to synthase expression reduced the magnitude of sensitivity to *γ* (Fig. S3). We also noted that both CL-RRD and CL-RD displayed less sensitivity to *γ* than CL-D and OL-D respecitvely, which indicates that additional feed-back to synthase expression can reduce sensitivity to dilution rate.

### Layered Feedback Decreases Sensitivity to Dilution

Previous studies have utilized forms of layered feedback to achieve robust regulation of molecules in biocircuits [4, 11]. We adapted their designs to design a simple layered feed-back controller in which AHLs acted through LuxR to directly repress LuxI via a LuxR repressable promoter and to indirectly repression LuxI through activating the expression of a secondary repressor that inhibits LuxI expression. We illustrate this architecture with both regulated and unregulated degradation in Fig. 3A, B, and we call these circuit CL-LRD (Closed Loop Layered, Repression Degradation) and CL-LRRD (Closed Loop Layered, Repression Regulated Degradation) respectively. The dynamics of LuxI under this regulation are modeled as follows:

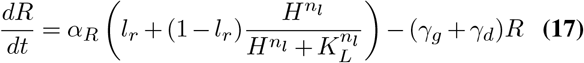

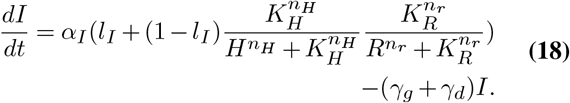

**Fig. 3.**
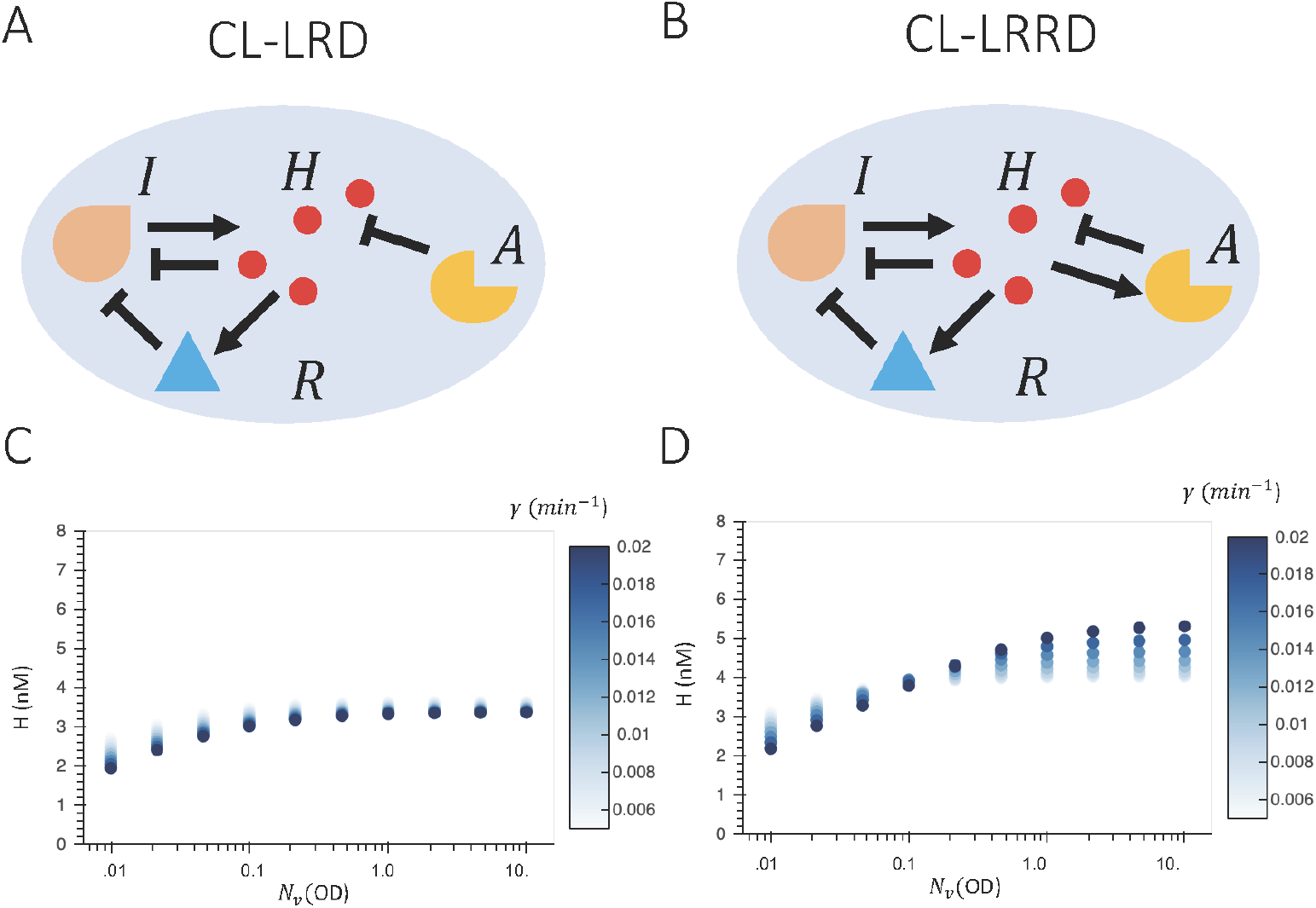
Diagrams (A,B) and steady state AHL levels (C, D) of layered feedback controller architectures. In panels A,B *H* represents AHLs, *I* respresents LuxI synthase, *A* represents AiiA degradase, and *R* represents secondary repressors. In panels C,D the steady state of AHLs (*H*) in nM is plotted for varying dilution rate(*γ* = *γ*_*r*_ = *γ*_*g*_) and OD (*N*_*v*_). Circuits were simulated for 1000 minutes, and the endpoint of the circuit was taken as the steady state of the circuit for a given set of parameters.

We found that for both layered architectures *H* was robust to variations in *N*_*v*_ with high *N*_*v*_, but CL-LRD displayed greater robustness to *γ* with with high *N*_*v*_ (Fig. 3B, C). We confirmed through sensitivity analysis that CL-LRD displayed reduced sensitivity to 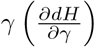 with high *N*_*v*_ S3G), but CL-LRRD did not (Fig. S4H). This suggests that layered feedback is most effective in regimes of strong degradase expression, which occurs when degradase expression is unregulated. CL-LRRD did not decrease sensitivity to *γ* much more than CL-RRD.

### Examination of Tunability of Dosage Control Circuits

Our above simulations used parameters such that the steady state of *H* was approximately 3-4 nM when *N*_*v*_ = 10, *γ* = 0.01min^−1^ for each circuit. We can tune parameters for each circuit to adjust the steady state output of *H*, but the control performance of the circuit depends on its parameters. To illustrate this, we characterized how the robustness of *H* in each circuit to varying *N*_*v*_ and *γ* depends on the strength of expression of the synthase and degradase. Since sensitivity to *N*_*v*_ and *γ* can only be calculated pointwise for each pair of *N*_*v*_, *γ*, we developed more concise metrics that summarized robustness to population size (*N*_*v*_) and dilution rate (*γ*) respectively. To describe robustness to population size we found the population density in which *H* was within 5% of the “plateau” value of *H*, or the value of *H* with large *N*_*v*_ (*N*_*v*_ = 10). We called this population density the controller population requirement, for it determines the smallest population density the controller needs to be robust to further increases in population density. Thus, a lower controller population requirement is desirable. For these calculations *γ* was held constant at *γ* = 0.001 min^−1^, but note that the controller population requirement may vary with *γ*. Since robustness to population size is achieved with high population size, we characterized robustness to *γ* in the limit of dense populations. To describe robustness to dilution rate (*γ*), we calculated the ratio 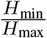 where *H*_max_ and *H*_min_ are the the maximum and minimum values of *H* achieved over a range of *γ* values when *N*_*v*_ = 10. We call this metric the *γ*_ratio_. In Fig. 4 we plot controller population requirements, *γ*_ratio_’s, and value of *H* when *N*_*v*_ = 10 and *γ* = 0.001 min ^−1^ for all circuits which incorporate degradases. We do not include circuits that do not have degradases, because these do not plateau at high *N*_*v*_.

**Fig. 4.**
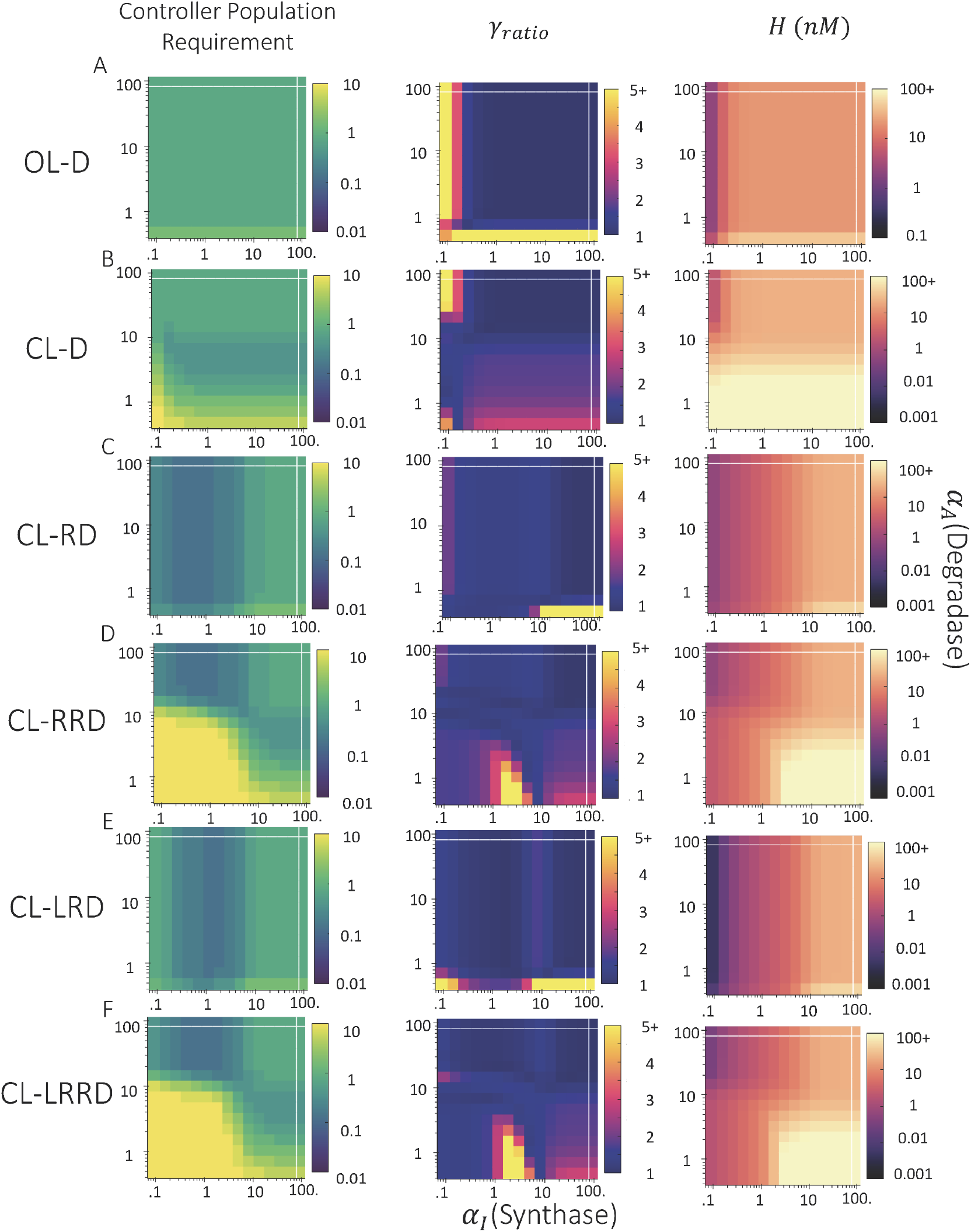
Performance of circuits with varied degradase (*α*_*A*_) and synthase (*α*_*I*_) promoter strengths. On each row from left to right are heat maps of *N*_*v*_ controller population requirements, *γ*_ratio_ and concentration of *H* for each simulated pair of *α*_*A*_, *α*_*I*_. The controller population requirements is defined as the population size (*N*_*v*_) where *H* is within 5% of the value of *H* with *N*_*v*_ = 10, and lower controller population requirements are desirable. For these simulations *γ* = 0.01 min^−1^. The *γ*_ratio_ is 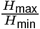 where *H*_max_,*H*_min_ are the maximum and minimum steady state values of *H* achieved at *N*_*v*_ = 10 for simulated values of *γ* ∈ [0.005, 0.02]. These are evaluated at *N*_*v*_ = 10 because our circuits in which *H* plateaus for high *N*_*v*_ plateau by *N*_*v*_ = 10. In the rightmost panel of each row is the value of *H* for *N*_*v*_ = 10, *γ* = 0.01min. In the *γ*_ratio_ panels the 5+ on the colorbar for *γ*_ratio_ indicates a regime where the *γ*_ratio_ > 5, but the same colorbar is used to allow more effective comparisons between circuits. Similarly, the 100+ on the colorbars for *H* indicates the regime where *H* > 100nM, but the same colorbar is used to allow more effective comparisons between circuits. For all values, circuits were simulated for 5000 min^−1^ and the end points were taken as the steady state of the circuit.

We found that circuits with unregulated synthase expression (CL-D, OL-D) displayed consistent controller population requirements of around *N*_*v*_ = 1 (Fig. 4A, B). Regulating synthase expression (Fig. 4C – F) yielded regimes of *α*_*A*_, *α*_*I*_ where the controller population requirement is below *N*_*v*_ = 1. However CL-RRD and CL-LRRD (circuits with regulated degradase and synthase expression) required high degradase promoter strengths *α*_*A*_ > 10 in order to maintain robustness to *N*_*v*_. With *α*_*A*_ > 10, these circuits often did not plateau with respect to *N*_*v*_. Additionally, both of these circuits had high *γ*_ratio_ for *α*_*A*_ > 10, which demonstrates that CL-RRD and CL-LRRD are not effective controllers with low *α*_*A*_. CL-RD and CL-LRD, the two closed loop controllers with constitutive degradase expression, displayed low *γ*_ratio_ and controller population requirenments for nearly all combinations of *α*_*A*_, *α*_*I*_. CL-LRD had a lower *γ*_ratio_ for the majority of (*α*_*A*_, *α*_*I*_), which suggests that the layered architecture reduces sensitivity to *γ*. Together, these results demonstrate that the control properties of all circuits are dependent on *α*_*I*_ and *α*_*A*_.

We next observed whether varying *α*_*I*_, *α*_*A*_ would change *H*, and thus make the circuit tunable. We noticed that tuning *α*_*I*_ had little effect on *H* for OL-D (Fig. 4 A, B), and that OL-D was restricted to a single value of *H* ≈ 20*nM* for high *α*_*A*_ and *α*_*I*_. We noted that when *α*_*A*_ and *α*_*I*_ are both high, we have that *I* ≫ *K*_*I*_ and *A* ≫ *K*_*A*_, so the dynamics of *H* become

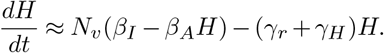

Thus the steady state of *H* is

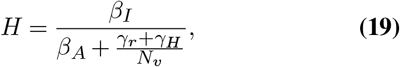

Note that when *N*_*v*_ → ∞ we have that 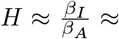 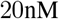 which is robust to *N*_*v*_ and both dilution rates. However, when this occurs the steady state of *H* is restricted to be 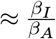. This scenario can be implemented using the OL-D architecture and appropriate values of *β*_*I*_ and *β*_*A*_. However, OL-D’s robustness to *γ* is lost when no longer working in the limit of saturated production and degradation.

On the other hand, the closed loop architectures were tunable. CL-D, CL-RRD CL-LRRD, could be tuned to reach *H* > 100nM, but at the cost of increased *γ*_ratio_ and higher controller population requirements (Fig. 4B, D, F). The highest concentration of *H* that circuits with constitutive degradase expression (CL-RD, CL-LRD) could reach without significant loss to robustness was 20nM (Fig. 4C, E).

### Examination of Properties of Circuit Trajectories

In addition to having robustness to dilution rate and *N*_*v*_, an ideal dosage control circuit should reach its steady state quickly and with minimal overshoot. In the context of dosage to a patient, an overshoot is undesirable for it would give a patient a potentially dangerously high dose of a drug. If a circuit takes too long to settle, then a patient could receive an inappropriate dose for a long time.

We first simulated example trajectories of each circuit where we held *γ*_*r*_ = *γ*_*g*_ = 0.01 min^−1^ and varied *N*_*v*_ (Fig. 5). In our simulations, all species were set to 0 initially. We first noticed that in all circuits (except OL) when *N*_*v*_ was high there was a significant overshoot. This is a property that can result from negative autoregulation [21] due to the delay in production of molecules that implement negative feed-back. Since it is undesirable for the circuits to have a significant overshoot, we quantified the overshoot of each circuit as 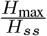 where *H*_max_ is the maximum concentration of AHLs reached before the system settles and *H*_*e*_ is the concentration of *H* the system settles to (Fig. 5, S3). First we noticed that circuits with more regulated species tended to have larger overshoots. For example, we noted that regulating degradase expression in CL-D increased the overshoot compared to OL-D (Fig. S4B, D), and that regulating synthase expression also increased overshoot compared to unregulated synthase expression. We also noticed that the over-shoot increased with *N*_*v*_ for all circuits but OL and OL-D (both of which displayed minimal overshoot) (Fig. S4). For circuits where the overshoot depended on dilution rate, the overshoot magnitude increased with decreasing dilution rates (*γ*_*g*_ = *γ*_*r*_) (Fig. S4 E - H). Unfortunately we noticed that CL-LRD, which has the best robustness to *γ*, has a much higher overshoot than the non layered circuits (Fig. 4G).

**Fig. 5.**
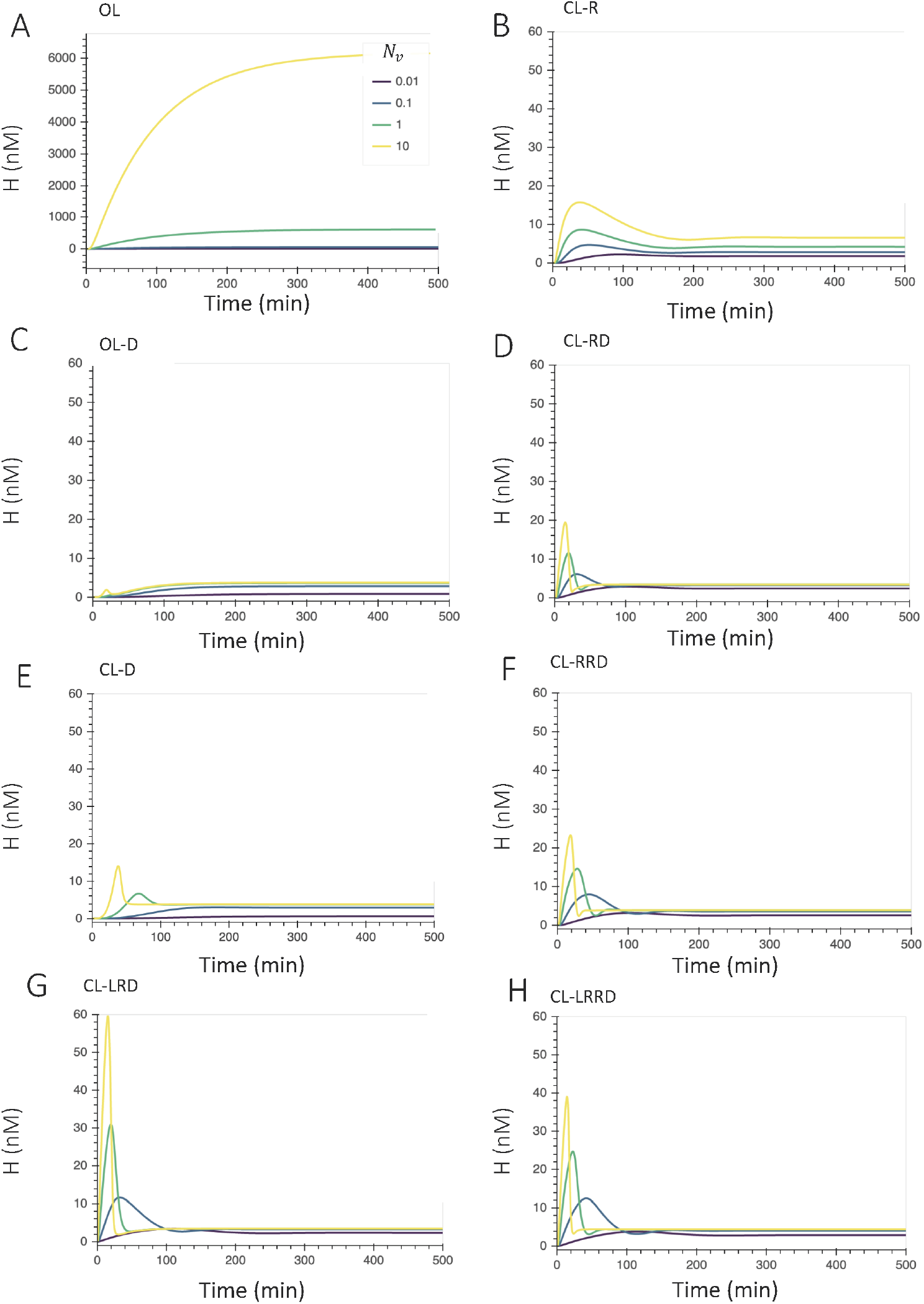
Open loop (A, C) and closed loop (B,D,E,F, G, H) example trajectories. For each circuit five trajectories with varying ODs (*N*_*v*_) were simulated for 500 min with *γ*_*g*_ = *γ*_*r*_ = 0.01 min^−1^. The legend in panel A applies to all figures. Other parameters governing these simulations can be found in Supp. Note 1.

We then characterized the 1% settling time for each of the circuits (Fig. 6). The 1% s settling time is defined as the time it takes for the circuit to reach a point within 1% its steady state value and cease changing beyond 1% of the end point. We note that our simulations are consistent with what was found in [15], where the time for CL-D decreased with *N*_*v*_ for fixed *γ*, and the settling time for the fully open loop circuit with no degradases does not vary with *N*_*v*_. We noticed that the settling time dynamics with respect to *N*_*v*_ and dilution rate seemed to cluster into three groups based on their mode of regulating degradases. The settling times for circuits which lacked degradases (OL and CL-R) displayed a strong dependence on dilution rate (Fig. 6A, B). In the limit of large *N*_*v*_ the settling times for circuits with unregulated degradase expression (OL-D, CL-RD, CL-LRD) became to robust to *N*_*v*_ (Fig. 6C, D, G). For the circuits with regulated degradase expression (CL-D, CL-RRD, CL-LRRD), settling times did not plateau with high *N*_*v*_ (Fig. 6E, F, H). We also found that circuits with regulated synthase expression had faster settling times than circuits with identical degradase regulation but unregulated synthase expression. Combined these results demonstrate degradases govern the settling times’ dependence on *γ*, *N*_*v*_, and synthase regulation can be used to speed up settling. Additionally, we characterized how simulations responded to a pulse AHLs into the bioreactor, similar to experiments performed in [15] (Fig. S5). We simulated each circuit for 1000 min, and then added a pulse of +*f* [*H*] where [*H*] is the concentration of AHLs at the end of the initial 1000 min simulation and *f* = .5. The post stimulus settling times were shorter than the settling times starting from all zero initial conditions. This makes sense, for much of the regulatory species are already in place at the time of the stimulus. Many of the behaviors for the settling times held for the post stimulus settling times. However, we noticed that the settling times for OL-D varied with *γ*, but the post stimulus settling times displayed robustness to *γ* in the limit of large *N*_*v*_, similar to those of CL-RD (Fig. S5C).

**Fig. 6.**
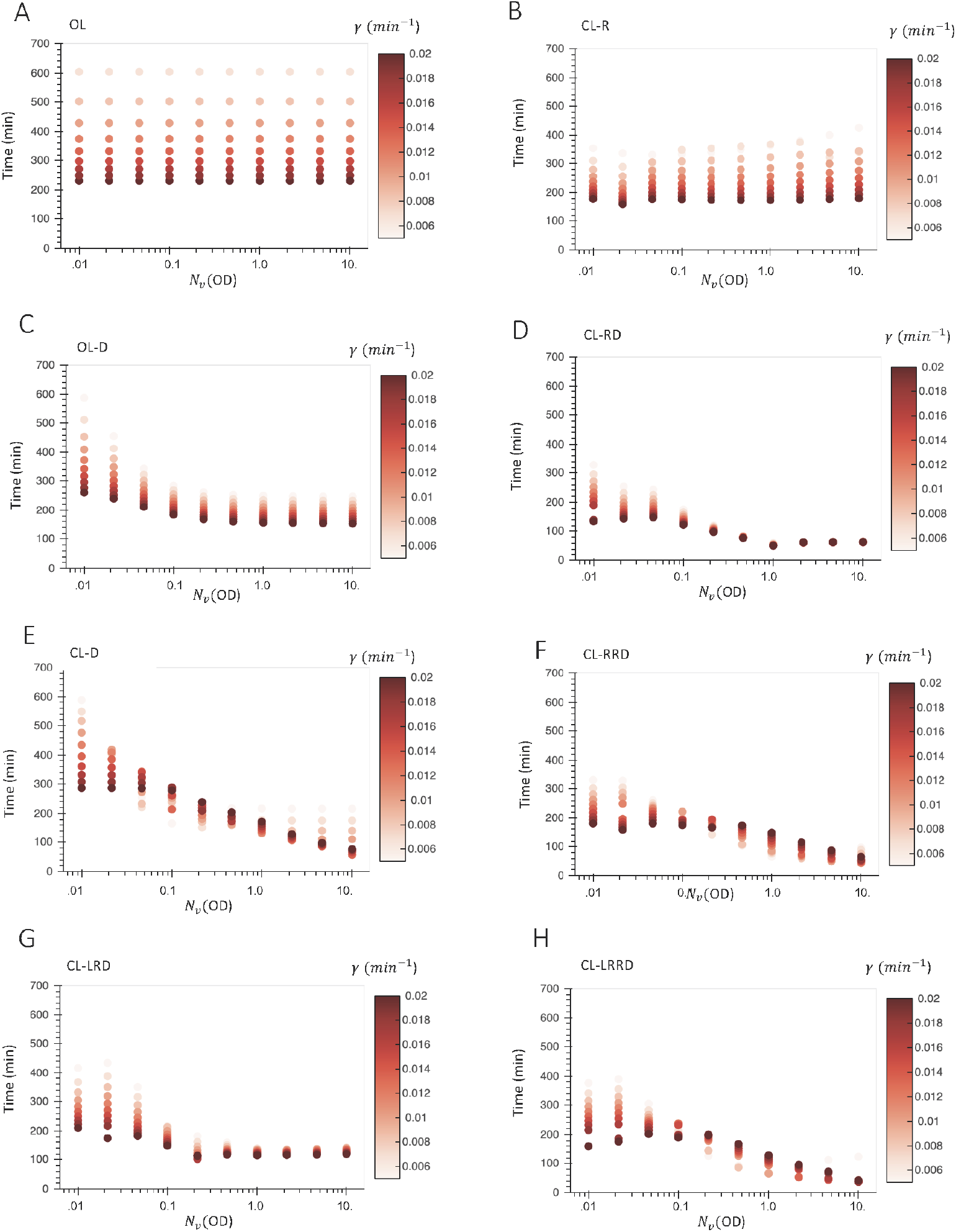
Settling times for varying dilution rate(*γ* = *γ*_*r*_ = *γ*_*g*_) and population sizes (*N*_*v*_) for various dosage control architectures. Settling time is defined as the time until *H* reaches and is maintained within 1% of its end point value during a 1000 min simulation. For these simulations all species were set to zero initially.

## Discussion

We explored several simple circuit architectures implementing dosage control. We used simplified analyses to gain intuition in which architectures could implement dosage control, and then simulated these circuits using biologically realistic parameters. Regulating AHL degradation and production rate are both forms of negative feedback that have potential to be utilized to create a circuit which implements population level control over a shared signal. We demonstrate that incorporating sufficiently strong degradation using AiiA molecules can yield robustness to population density within high population densities. The circuits with greatest robustness to dilution rates also regulated *H* through controlling synthase expression, and we found that a layered repression could yield greater robustness to dilution rates in some cases. We also evaluated how robustness was dependent on tunable parameters in each circuit, and we examined properties of the trajectories of circuits.

All of our experiments were performed *in silico* using biologically realistic parameters, most of which come from experimental data [15, 20]. However, some of these parameters are likely to be incorrect under different contexts, such as varying growth rates and reactor conditions. Furthermore, our *in silico* assumptions may not be valid in certain *in vivo* environments. For example, our current model of AiiA degradation assumes that degradation rate is proportional to *H* and a Hill function of AiiA. This Hill function has quite high ultrasensitivity, which indicates that the dynamics of AiiA may not be accurately modeled by a Hill function. Additionally, there is likely some regime of AHL where degradation rate is no longer proportional to AHL, but we assume that we are dealing with small enough AHL concentrations that it is approximately linear. Additional *in silico* analysis should also focus on how different dosage control architectures perform when we change our assumptions about how the Lux quorum sensing system works. We thus need to characterize these circuit architectures more thoroughly through *in vivo* experiments. However, analysis from *in silico* experiments has yielded important insight into the necessary conditions for dosage control.

Although some of these circuts are able to implement dosage control, one drawback of all circuits is that their outputs are dependent on molecular parameters, such as maximum degradation rate of AHLs by AiiA and maximum production rate of AHLs by a synthase molecule. As mentioned previously, these parameters may not be constant under varied environmental conditions, such as different medias, which would make our predictions inaccurate. Additional future work should also focus on designing controllers which are robust to variations in parameters intrinsic to the circuit.

As mentioned earlier, one challenge in synthetic biology is to engineer microbial consortia that have behaviors and functions that are not present or possible in single cells [1, 2], and population controllers are needed to address this challenge. In our work we provide insight into circuit architectures for controlling a shared signal that is produced by a homogeneous population of cells. Future work should explore how these controllers could be utilized for other applications in synthetic biology.

## Acknowledgements

We would like to thank Xinying Ren, John Marken, William Poole, and the Murray lab for useful discussions and insights throughout the project. This research is supported in part by the Institute for Collaborative Biotechnologies through contract W911NF-19-D-0001 from the U.S. Army Research Office. The content of the information on this page does not necessarily reflect the position or the policy of the Government, and no official endorsement should be inferred.

## Code Availability

All code for figures and results of this study can be found at https://github.com/sophiejwalton/dosage_control.

---

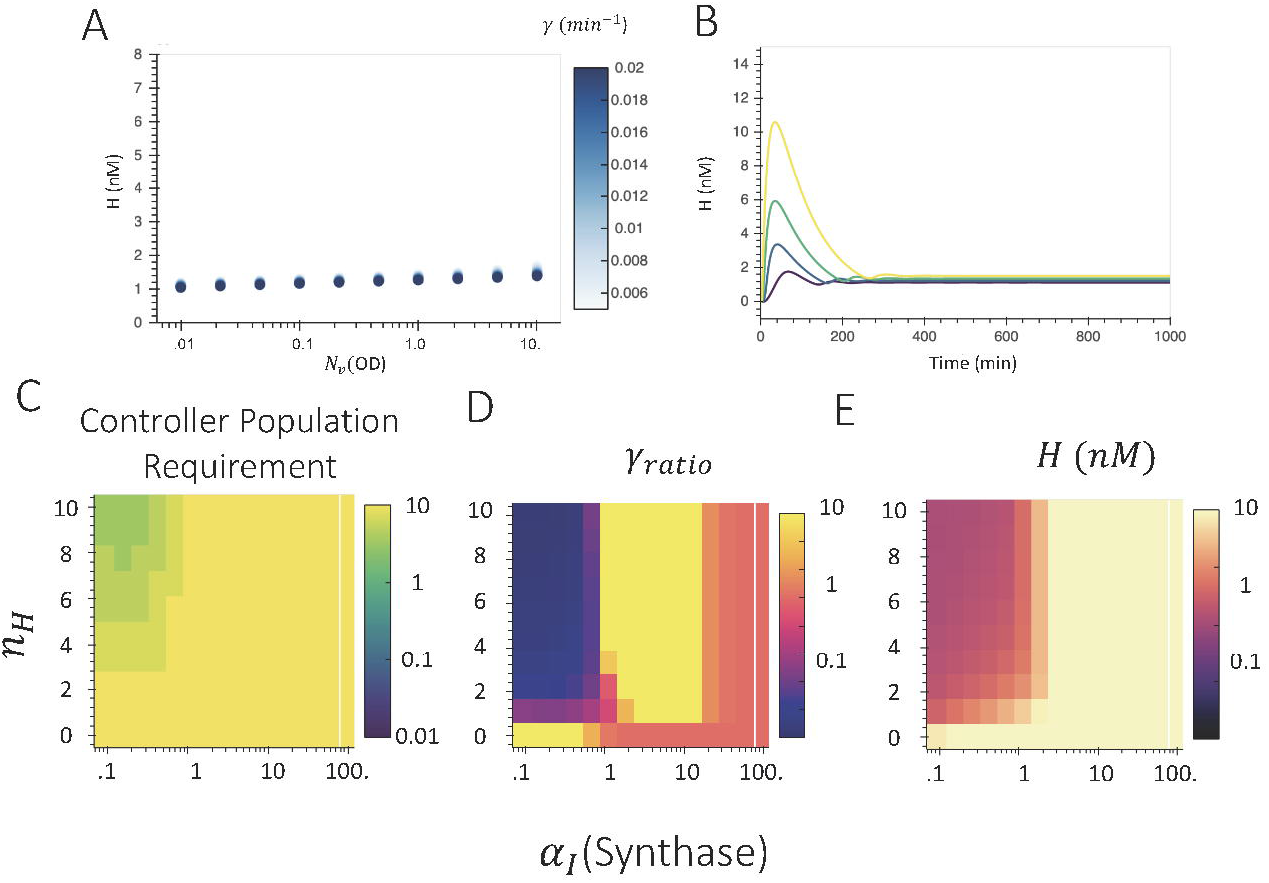

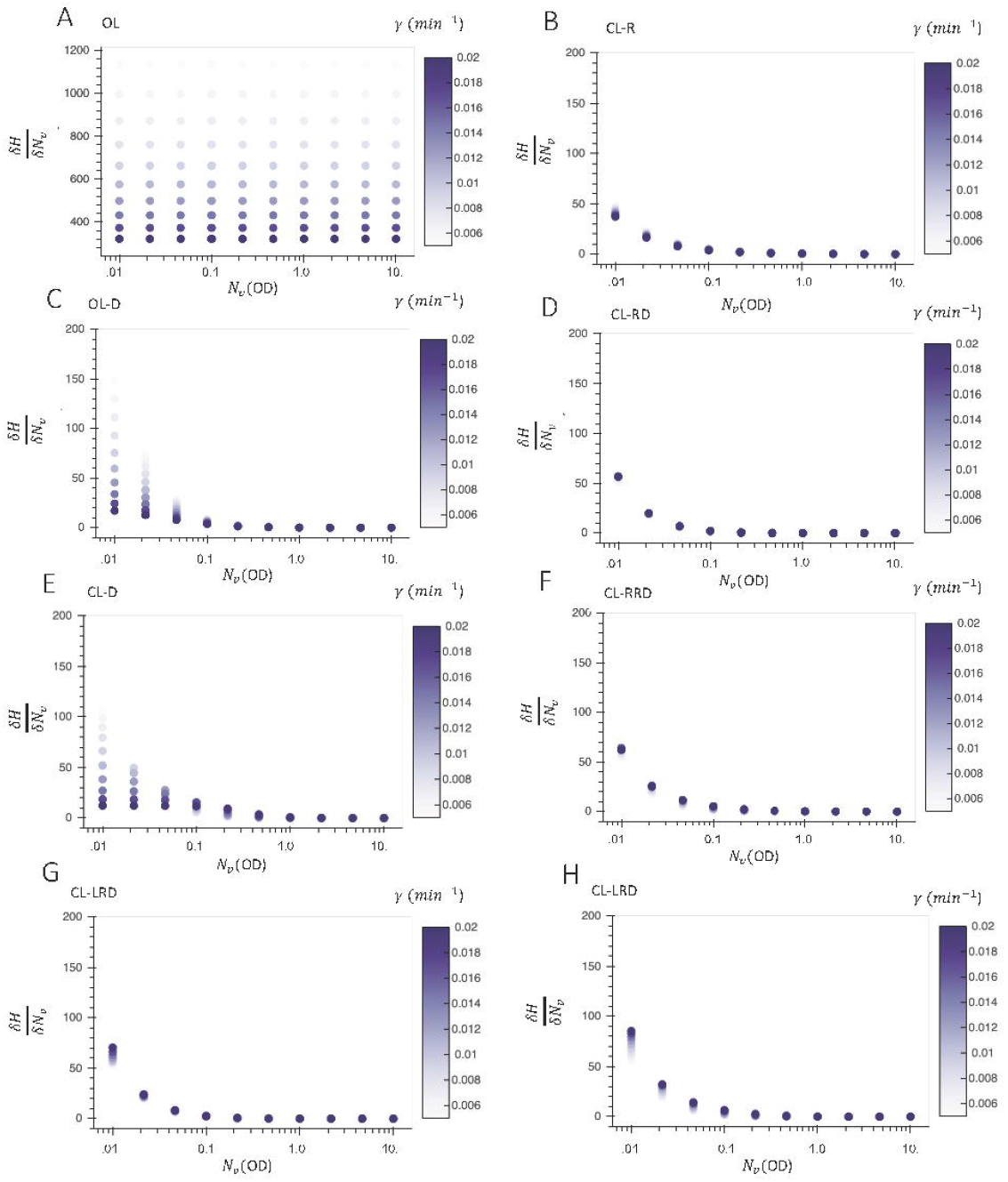

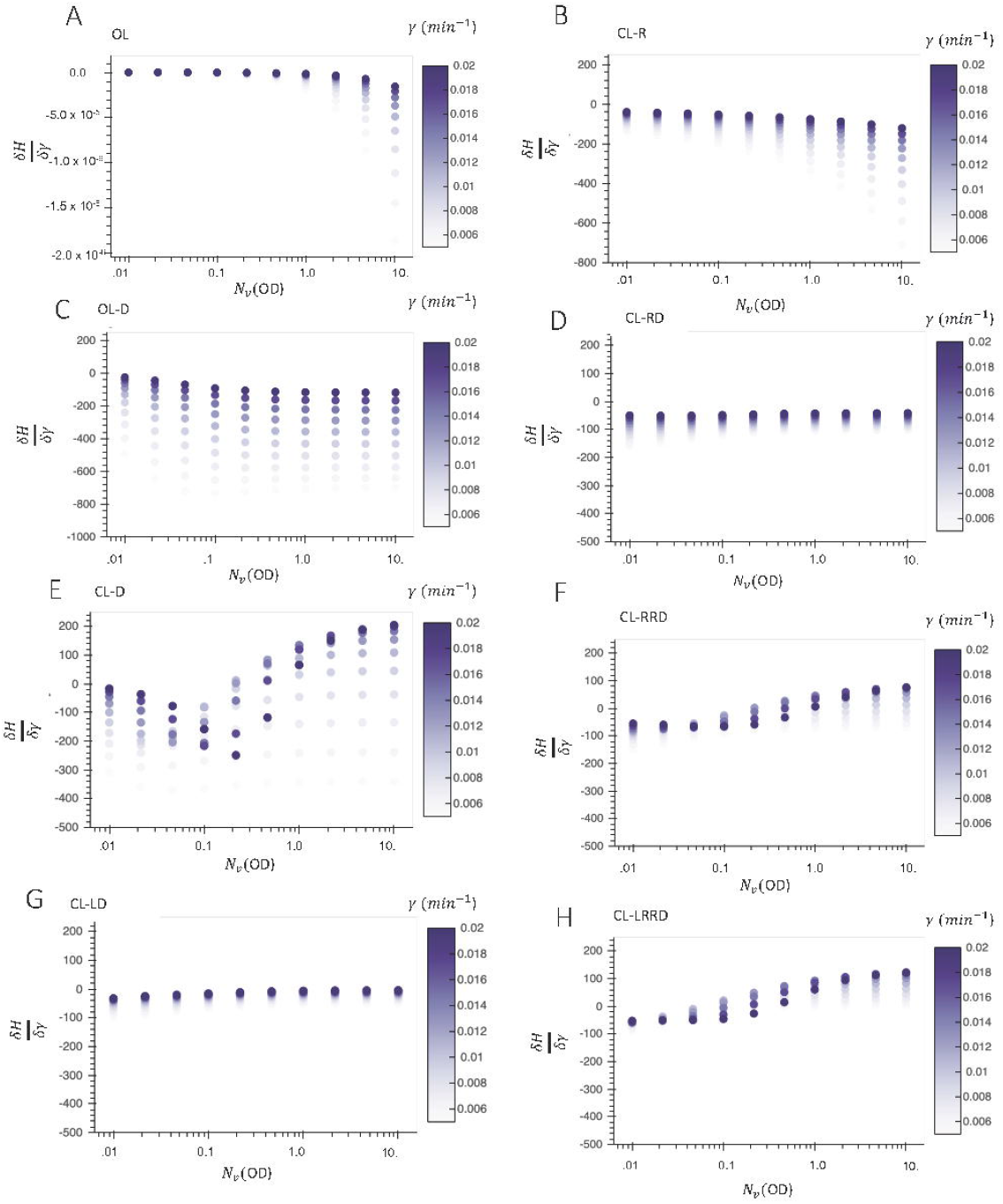

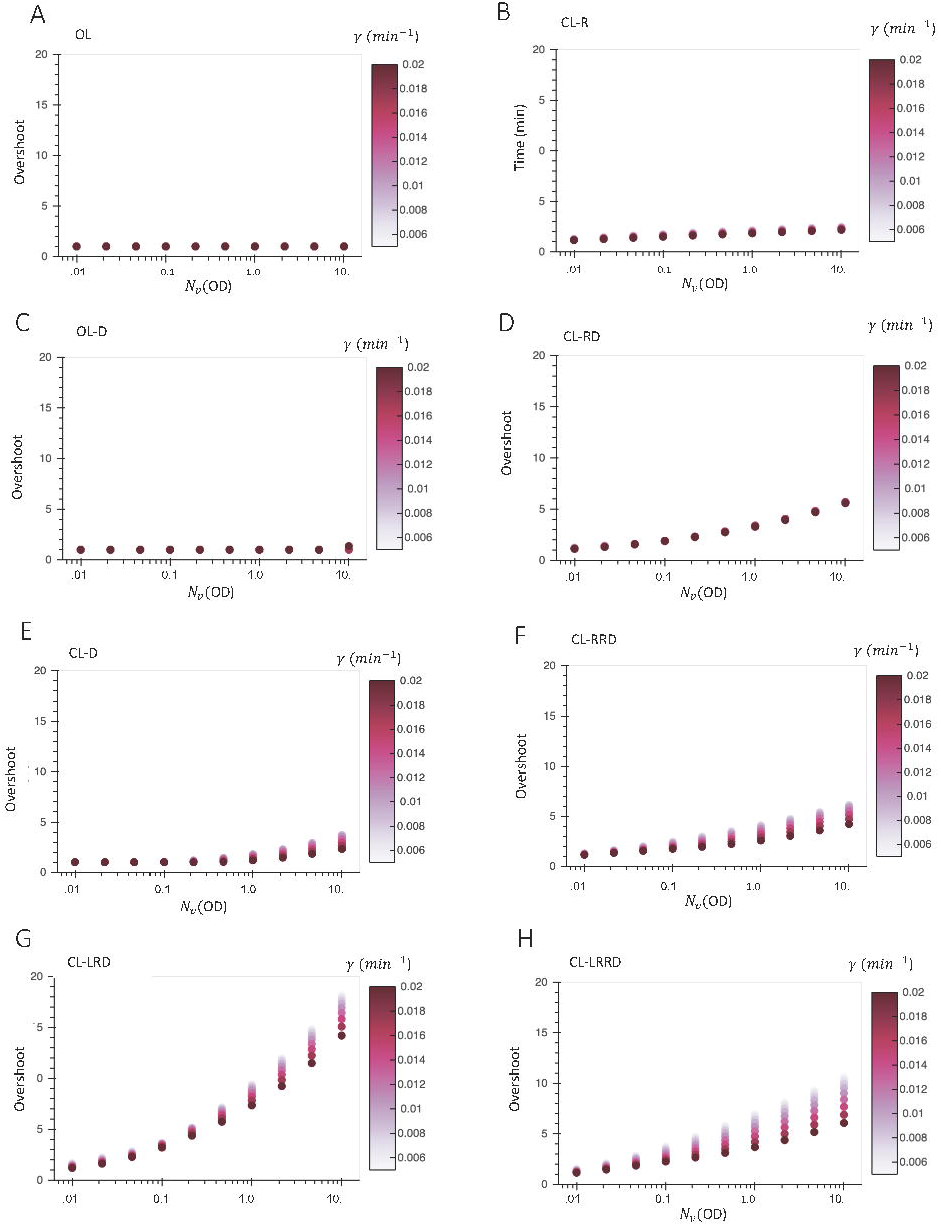

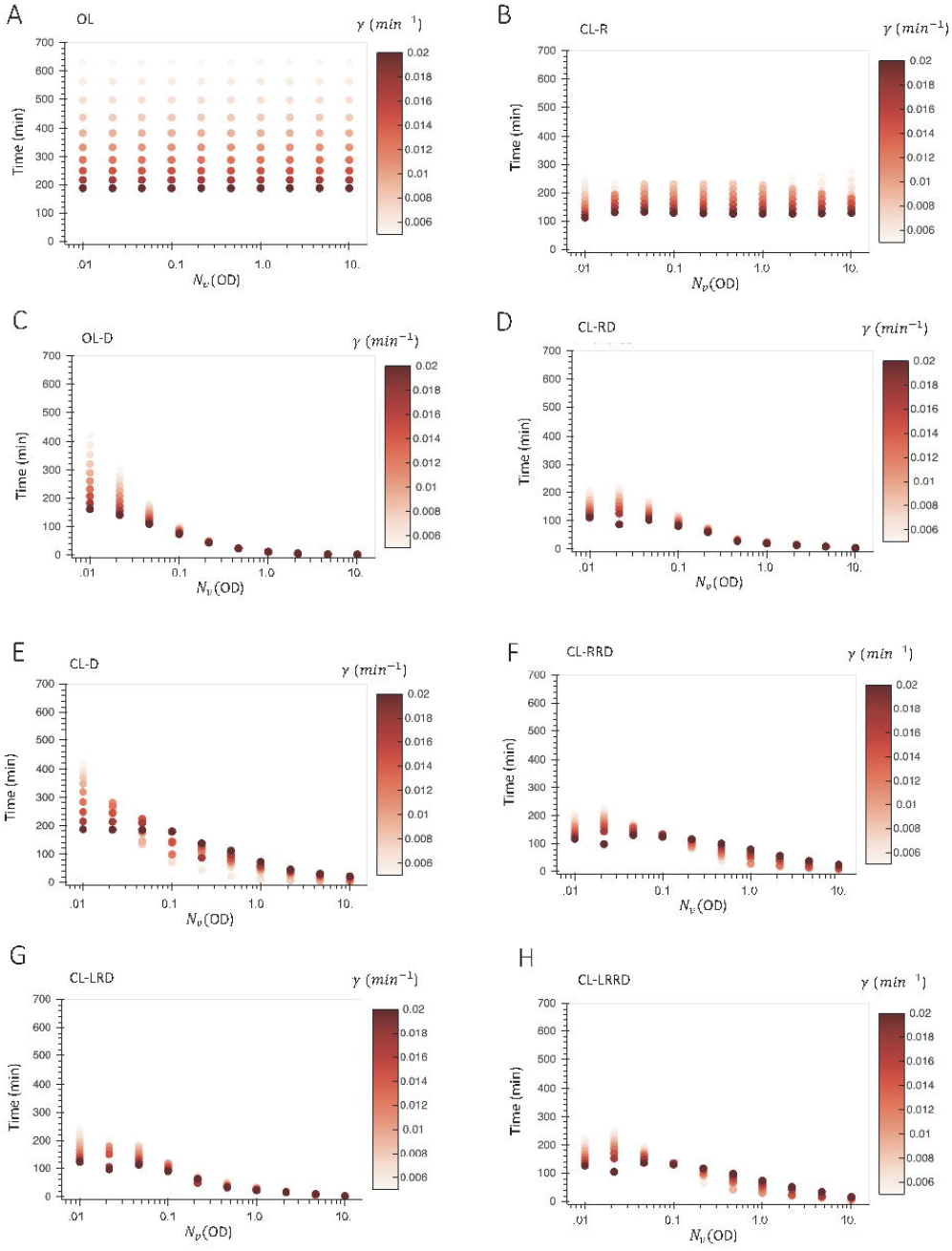

