## Supplemental Notes and Figures for "Analysis of Circuits for Dosage Control in Microbial Populations"

### Supplementary Note 1: Mathematical Modeling and Simulations

All simulations were performed using Python 3.7 and Scipy libraries. Values were estimated based on results from [20] and [15]. Unless otherwise indicated parameters used for each circuit can be found below:

| Parameter | Units | Estimated Value | Circuit |
|---|---|---|---|
| $\alpha_A$ | nM min$^{-1}$ | 1 | CL-RD, CL-LRD, OL-D |
| $\alpha_A$ | nM min$^{-1}$ | 15 | CL-D, CL-RRD, CL-LRRD, CL-LRRD |
| $\alpha_I$ | nM min$^{-1}$ | 1 | OL, CL-RD, CL-RRD |
| $\alpha_I$ | nM min$^{-1}$ | 2 | CL-LRRD |
| $\alpha_I$ | nM min$^{-1}$ | 3 | CL-LRD |
| $\alpha_I$ | nM min$^{-1}$ | .1 | CL-D |
| $\alpha_I$ | nM min$^{-1}$ | .12 | OL-D |
| $\alpha_I$ | nM min$^{-1}$ | .5 | CL-R |
| $\alpha_R$ | nM min$^{-1}$ | 2 | CL-LRD |
| $\alpha_R$ | nM min$^{-1}$ | 1 | CL-LRRD |
| $\gamma_d$ | min$^{-1}$ | 0.0173 | - |
| $\beta_I$ | nM min$^{-1}$ | 6.83 | - |
| $\beta_A$ | min$^{-1}$ | .33 | - |
| $l_I$ | - | 0.02 | - |
| $l_A$ | - | 0.027 | - |
| $l_R$ | - | 0.02 | - |
| $K_L$ | nM | 474 | - |
| $n_L$ | - | 1. | - |
| $K_H$ | nM | 1. | - |
| $n_H$ | - | 1.8 | - |
| $K_I$ | nM | 6.74 | - |
| $n_I$ | - | 3.51 | - |
| $K_A$ | nM | 18.7 | - |
| $n_A$ | - | 9.7 | - |
| $K_R$ | nM | 1 | - |
| $n_r$ | - | 2 | - |

Sensitivities present in Fig. S2 and Fig. S3 were calculated by taking the Jacobian of the differential equations used to model the circuit at the steady state circuit value for a given set of parameters. The numdifftools package (version 0.9.39) was used to compute Jacobians.

### Supplementary Figures

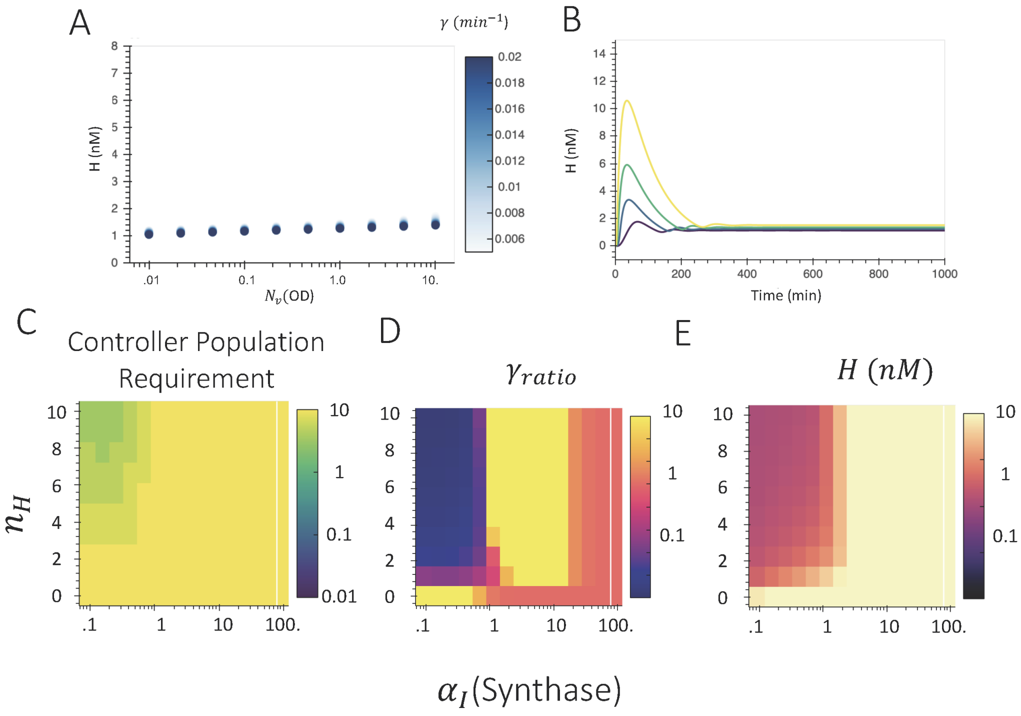


**Fig S1.** Evaluation of CL-R with large $n_H$. Example steady states (A) and trajectories (B) of CL-R with $n_H = 10$. All other parameters can be found in Supp. Note 1. Evaluation of Controller Population Requirement (C), $\gamma_{\text{ratio}}$ (D), and $H$ (E) for varied $n_H, \alpha_I$ in CL-R.

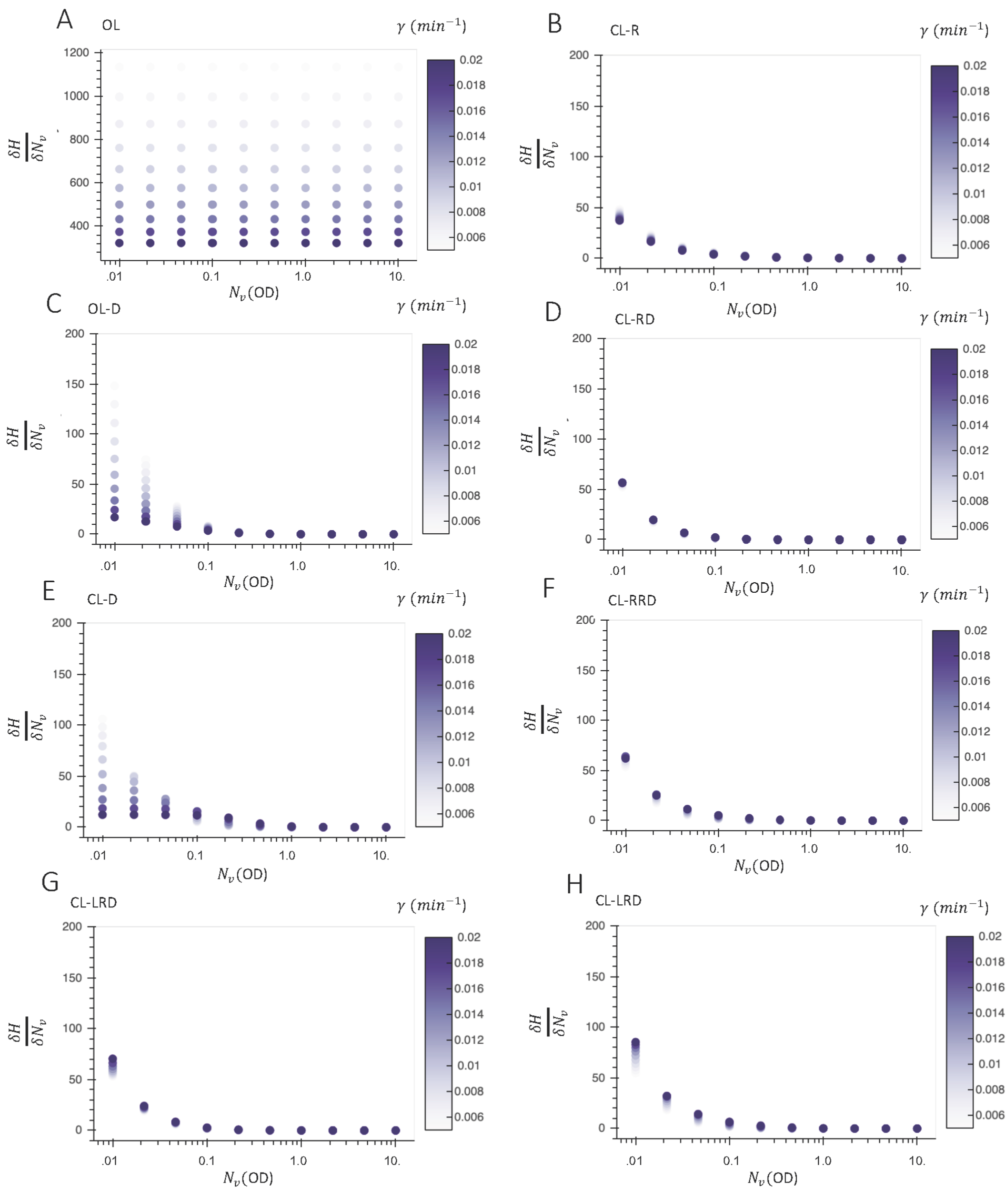


**Fig S2.** Sensitivity of $H$ to $N_v$ $\left(\frac{\partial H}{\partial N_v}\right)$ for varying dilution rate($\gamma = \gamma_r = \gamma_g$) and OD $(N_v)$ for various dosage control architectures.

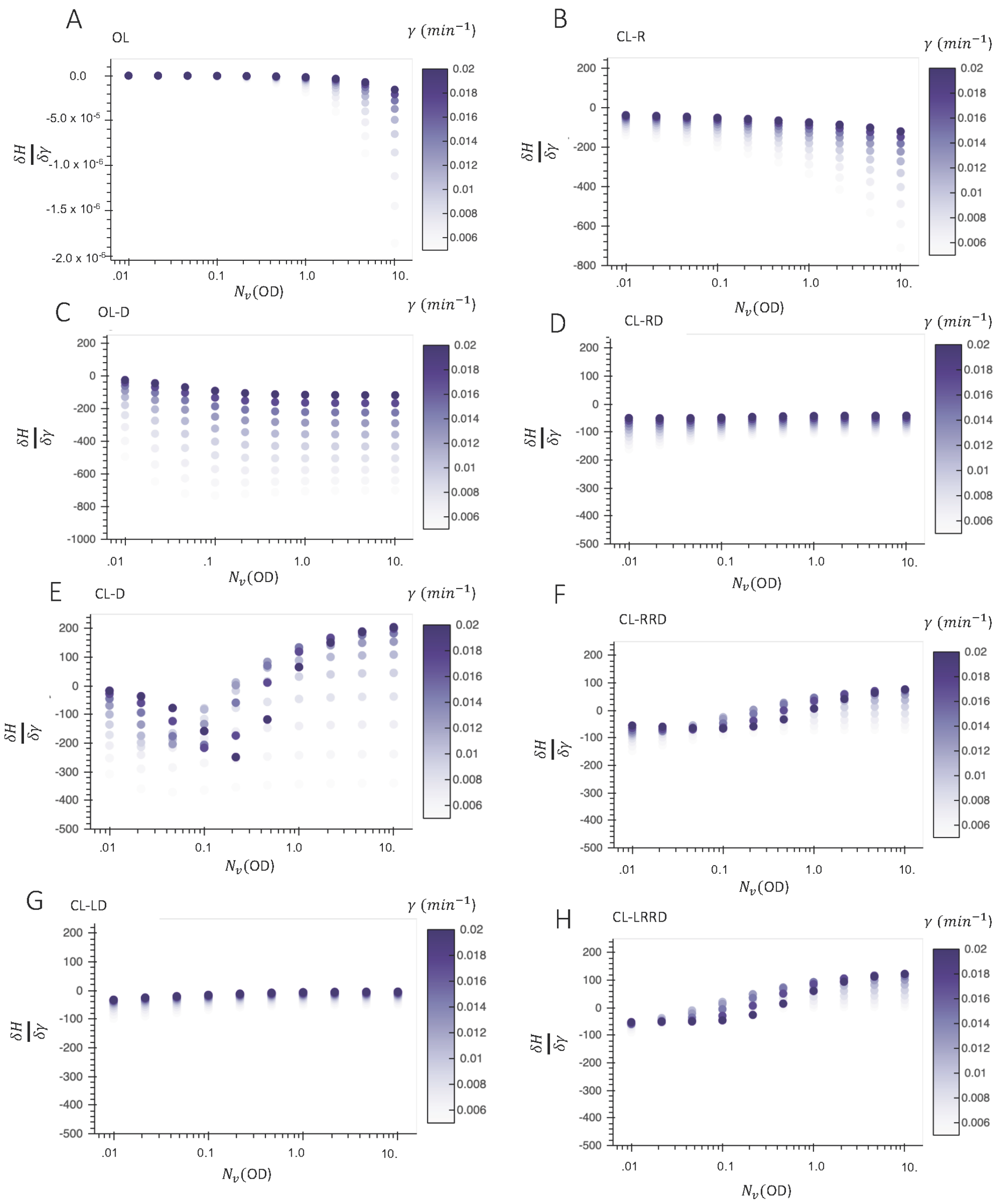


**Fig S3.** Sensitivity of $H$ to $\gamma$ $\left(\frac{\partial H}{\partial \gamma}\right)$ for varying dilution rate($\gamma = \gamma_r = \gamma_g$) and OD $(N_v)$ for various dosage control architectures.

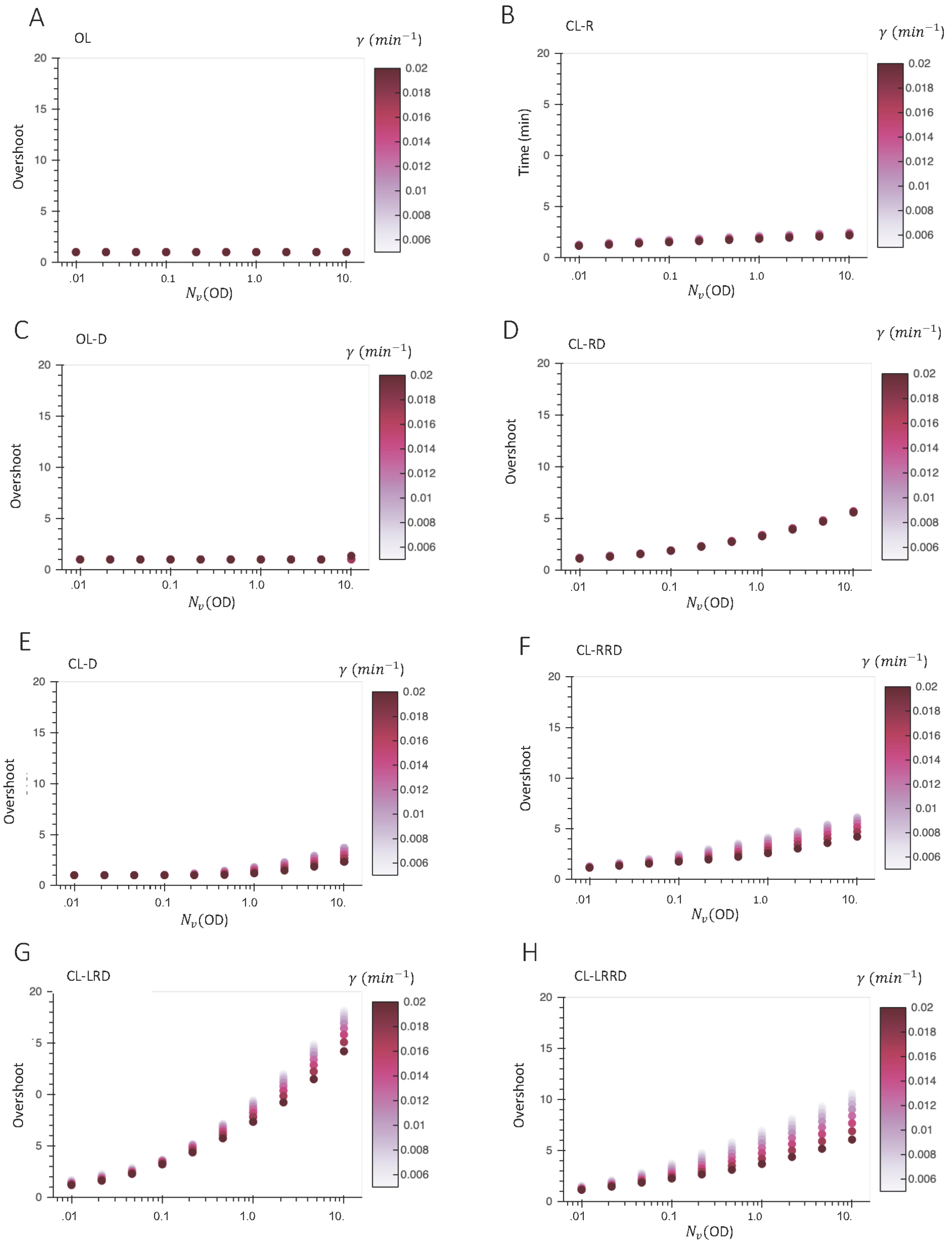


**Fig. S4** AHL ($H$) overshoot for varying dilution rate($\gamma = \gamma_r = \gamma_g$) and OD ($N_v$) for various dosage control architectures. The overshoot is defined as $\frac{H_{\text{max}}}{H_e}$ where $H_e$ is the value of $H$ at the end of the simulation and $H_{\text{max}}$ is the maximum value of $H$ achieved during the stimulation. Circuits were simulated for $1000$ minutes, and the endpoint of the circuit was taken as the steady state of the circuit for a given set of parameters.

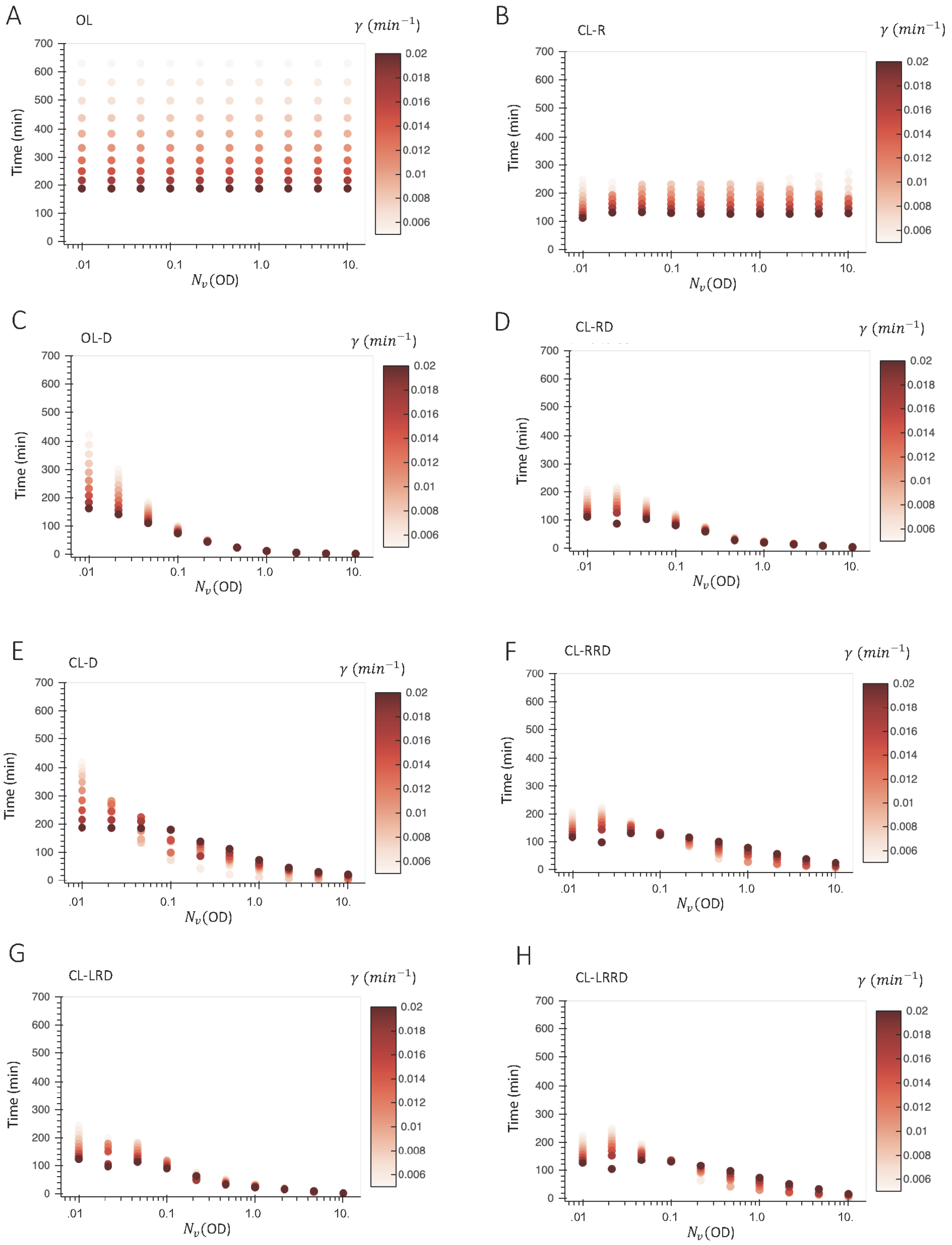


**Fig S5.** Settling times (time to reach and maintain within $1\%$ of steady state $H$) after a pulse of $H$ for varying $\gamma = \gamma_g = \gamma_r$ and $N_v$. All circuits were simulated for $1000$min starting from all zero initial conditions. After $1000$ min a pulse of $+.5$ AHL ($H$) was added, where $H$ is the concentration of AHLs in nM at the end of the initial $1000$ min simulation. The circuits were simulated for another $1000$min to find the settling time after the stimulus.
